# *Kras* loss of heterozygosity promotes MAPK dependent pancreatic ductal adenocarcinoma and induces therapeutic sensitivity

**DOI:** 10.1101/2023.08.29.552598

**Authors:** Sigrid K. Fey, Arafath K. Najumudeen, Catriona Ford, Kathryn Gilroy, Rosie Upstill-Goddard, David K. Chang, William Clark, Colin Nixon, Simon T. Barry, Jennifer P. Morton, Andrew D. Campbell, Owen J. Sansom

## Abstract

Pancreatic cancer is characterised by the prevalence of oncogenic mutations in *KRAS*. Previous studies have reported that altered *Kras* gene dosage drives progression and metastatic incidence in pancreatic cancer. While the role of oncogenic *KRAS* mutation is well characterised, the relevance of the partnering wild-type *KRAS* allele in pancreatic cancer is less well understood and controversial. Using *in vivo* mouse modelling of pancreatic cancer, we demonstrate that wild-type *Kras* restrains the oncogenic impact of mutant *Kras*, and drastically impacts both *Kras*-mediated tumourigenesis and therapeutic response. Mechanistically, deletion of wild-type *Kras* increases oncogenic *Kras* signalling through the downstream MAPK effector pathway, driving pancreatic intraepithelial neoplasia (PanIN) initiation. In addition, in the KPC mouse model, a more aggressive model of pancreatic cancer, loss of wild-type KRAS leads to accelerated initiation but delayed tumour progression. These tumours had altered stroma, downregulated *Myc* levels and an enrichment for immunogenic gene signatures. Importantly, loss of wild-type *Kras* sensitises *Kras* mutant tumours to MEK1/2 inhibition though tumours eventually become resistant and then rapidly progress. This study demonstrates the repressive role of wild-type *Kras* during pancreatic tumourigenesis and highlights the critical impact of the presence of wild-type KRAS on tumourigenesis and therapeutic response in pancreatic cancer.

## Introduction

Oncogenic *KRAS* mutations are common in cancer, and almost ubiquitous in pancreatic cancer, with mutations detected in around 95 % of patients. This significant prevalence of *KRAS* mutations is compounded by the relative lack of success in direct therapeutic targeting of oncogenic KRAS. There have been recent successes in this area, with the development and subsequent clinical use, albeit in different indications, of covalent binding inhibitors of KRAS-G12C such as AMG-510 (Sotorasib) ^1^. While many years of research have driven our understanding of the biochemical, cellular and indeed tissue level impact of oncogenic *KRAS* mutations in cancer, there is now growing evidence that not only mutation, but also allelic balance at oncogenic loci such as *KRAS*, and the resulting oncogene dosage, are critical modifiers of oncogenic signalling, tumour progression, and ultimately patient prognosis and therapeutic efficacy. It is noteworthy that this is not restricted to pancreatic cancer, with similar oncogene allele imbalances detected in cancers of different tissues of origin such as lung, colorectal, melanoma, breast and ovarian cancers ^2–7^. Moreover, copy number variants are not only limited to mutated oncogenic *KRAS* but are also detected with different driver genes (*NRAS*, *BRAF, EGFR* and *MYC*) ^3,4,6–9^.

In the case of *KRAS*, there is now clear evidence that alterations in allele frequencies or gene dosage at oncogene loci can have a dramatic impact upon tumourigenesis and tumour-associated phenotypes across tumour types ^2,6,10^. *KRAS* allelic imbalances are frequently observed (55 %) in mutant *KRAS* driven cancer across many tissues of origin ^5^. The effect of *KRAS* allelic imbalance upon therapeutic efficacy was observed in a study by Burgess et al., where it was demonstrated that the relative balance between mutant and wild-type *KRAS* alleles determined clonal fitness and sensitivity to MAPK targeting therapeutic approaches in acute myeloid leukemias (AML)^11^. Allelic imbalances at the *KRAS* locus have previously been reported in pancreatic ductal adenocarcinoma (PDAC), the most common type of pancreatic cancer, and were associated with early tumour progression ^2^. There is evidence that *KRAS* gene dosage has impact on PDAC evolution, indicated by variations in cell morphology, histopathology, and clinical outcome in patients. Moreover, copy number alteration of *KRAS*, such as copy number gain of oncogenic *KRAS*, loss of heterozygosity (LOH) or allelic imbalance can be beneficial for tumour growth. In pancreatic cancer increased *KRAS* gene dosage is associated with early tumourigenesis, progression and increased incidence of metastasis *in vivo* ^2^.

There is growing evidence that *KRAS* allelic imbalance might significantly affect mutant KRAS signalling through increased oncogenic *KRAS* gene dosage relative to the wild-type allele ^2,12^. This suggests that there are three distinct types of human KRAS mutant cancers, tumours that contain the wild-type *KRAS* allele and are heterozygous for *KRAS*, tumours where the mutant *KRAS* allele is exclusively expressed (homozygous), and tumours that have mutant *KRAS* copy number gain ^12^. This could conceivably impact tumour initiation, and progression, and may even modify responses to therapy through enabling intrinsic and acquired resistance.

We demonstrate the functional relevance of wild-type KRAS and understand mechanisms which govern the role of wild-type KRAS in the processes of tumour initiation and progression in pancreatic cancer. We use genetically engineered mouse (GEM) models of pancreatic cancer, allowing deletion of wild-type *Kras*, in combination with preclinical trialling of clinically relevant target therapeutic approaches to carefully dissect each of these features. We demonstrate that the presence of wild-type KRAS restrains mutant KRAS signalling and wild-type KRAS loss results in increased tumour PanIN *in vivo*. Importantly, we show that loss of wild-type *Kras* alters the progression of pancreatic tumours and leads to increased immune cell infiltration in preclinical models of pancreatic cancer. Using late-stage treatment intervention studies we show that loss of wild-type *Kras* enhances sensitivity to MEK inhibition *in vivo*. This study highlights the impact of *Kras* allelic imbalance and the critical role of wild-type KRAS in initiation, progression and therapeutic response in pancreatic cancer.

## Materials and Methods

### ICGC/APGI RNAseq patient dataset

Transcriptomic profiling and molecular characterisation of the ICGC PACA-AU cohort was performed as previously described (Bailey et al.). Copy number calls for samples included in the PCAWG project were obtained from ICGC. Samples were defined as either KRAS balanced or KRAS imbalanced. Samples classed as KRAS balanced had an equal number of copies of the major allele and minor alleles, while samples classed as KRAS imbalanced had more copies of the major allele compared to the minor allele.

### Mouse models and experiments

Experiments using Genetic Engineered Mouse Models (GEMMs) were performed according to the UK Home Office regulations (licenses 70/8646 and PP3908577), under approval by the animal welfare and ethical review board (AWERB) of the University of Glasgow.

The *in vivo* studies were performed using the KPC mouse model ^13^. The alleles were used as follows - *Pdx-1*-*Cre* ^14^, lox-stop-lox-*Kras*^G12D^ ^15^, lox-stop-lox-*Trp53*^R172H^ ^16^. *Kras*^fl^ was generated by the International Mouse Phenotyping Consortium (IMPC). The smallest sample size that could give a significant difference was chosen (specified in relevant figure legends), in accordance with the 3Rs. No formal randomization was used, and the experimenter was blinded to genotypes.

Mice were bred on a mixed background. Mice of both sexes were used for tumour studies and monitored at least three times weekly and sampled when exhibiting clinical signs of PDAC (abdominal swelling, jaundice, hunching, piloerection, or loss of body conditioning).

### Tumour monitoring

High-resolution ultrasound imaging was performed using a Vevo3100 System (with MX550D (25-55mHz) transducer (VisualSonics). To identify tumour-bearing animals, mice were assessed twice weekly for palpable abdominal masses, and where abdominal mass was present, pancreatic tumour burden was confirmed, and the size of the tumour at the enrolment was measured by high-resolution ultrasound imaging. Tumour growth was then monitored once weekly by ultrasound imaging until reaching clinical endpoint using VeVo Lab (Fujifilm, Visual Sonics, VeVo LAB 1.7.1, Toronto, ON, Canada).

### *In vivo* treatment experiments

For drug treatment studies, pancreatic malignancy was confirmed by abdominal palpation and ultrasound imaging. Mice were randomly assigned to cohorts. Mice were treated from palpable pancreatic tumour burden until reaching clinical endpoint or from indicated timepoints (as indicated in figure legends). The MEK inhibitor (AZD6244 or selumetinib) was administered in a concentration of 25 mg/kg twice daily via oral gavage in a vehicle of 0.5% HPMC and 0.1% Tween-80.

### Pancreatic intraepithelial neoplasia (PanIN) scoring

For pancreatic intraepithelial neoplasia (PanIN) scoring mice were sampled at indicated time points and PanINs were scored from whole H&E sections and normalized to mm^2^ pancreas of the whole section. PanINs were scored according to John Hopkins University Department of Pathology classification of PanIN ^17,18^.

### Immunohistochemistry

IHC was performed on formalin-fixed pancreatic sections using standard protocols. Haematoxylin-and-eosin (H&E) staining was performed using standard protocols. Picro Sirius Red staining technique was used to stain collagen within tissue sections as described previously ^19^. Primary antibodies used for immunohistochemistry were as follows: phospho-ERK1/2 (1:400, Cell Signaling Technologies #9101), cMYC (1:200 Abcam #ab32072), p53 (1:100, Cell Signaling Technologies #2524), αSMA (1:25000, Sigma-Aldrich #A2547), p21 (1:150, abcam #107099), Ki67 (1:100, ThermoFisher #RM-9106), Tenascin C (1:300, Sigma-Aldrich #T3413), Dusp6 (1:250, abcam #Ab76310). *In situ* hybridisation (ISH) (RNAscope) was performed according to the manufacturer’s protocol (Advanced Cell Diagnostics RNAscope 2.0 High Definition–Brown) for *Dusp5 (*Bio-Techne #475698), *Dusp6 (*Bio-Techne #429321) and *cmyc (*Bio-Techne #508808). Sections were counterstained with hematoxylin and covered by a coverslip using DPX mountant (CellPath, UK).

### Assaying proliferation *in vivo*

Proliferation levels were assessed by measuring BrdU incorporation. Mice were injected intraperitoneally with 250µl of BrdU (Amersham Biosciences) 2 hours before being sacrificed. IHC staining for BrdU was then performed using an anti-BrdU antibody (1:200, BD Biosciences #347580).

### Western Blotting

Snap frozen PDAC tissue was homogenized in RIPA lysis buffer (Tris pH 7.4, NaCl, Triton X-100, SDS, dH_2_O) using a Precellys homogenizer. Lysates were centrifuged for 10 min at full speed. Protein concentration in the supernatant was determined using a BCA assay kit (Thermo scientific # 23225). Samples were boiled in loading buffer for 5 min at 95 °C and allowed to cool down to room temperature prior to loading, 20ug protein were separated on a 4-12% gradient pre-cast gel (Novex #NP0322PK2) and run in MOPS running buffer. Samples were transferred onto PVDF membrane (Millipore, #IPFL00010). Membrane was blocked in 5 % BSA (Sigma #A9647) in TBS-T for 1 h at room temperature. Primary antibody was incubated in 5 % BSA in TBS-T or 5 % milk in TBS-T at 4°C overnight. Primary antibodies were used as follows: (KRASG12D 1:1000, CST #14429; ß-actin 1:2000, SIGMA #A2228; ERK1/2 1:1000, CST #4695; pERK1/2 1:1000, CST #9101; MEK1/2 1:1000, CST #8727; pMEK1/2 1:1000, CST #2338; AKT 1:1000, CST #9272; pAKT (Ser473) 1:2000, CST #4060; PTEN 1:1000, CST #9559; DUSP6 1:1000, abcam #ab76310). Secondary antibody (α-rabbit Dako #P0448, α-mouse Dako #P0447) was diluted 1:2000 in 5% BSA/TBS-T and incubated for 1h at RT. Membranes were incubated with ECL plus working reagent (Thermo Scientific catalogue no. 32132) for 2 min and bands were detected using a Chemidoc Imager (Biorad). Membranes were briefly washed with TBS-T and stripped with Replot Plus (Millipore #2504) for 10 min at RT and rinsed with TBS-T.

### RAF-RBD assay (Ras Pull-down activation assay)

Snap frozen PDAC tissue was homogenized in lysis buffer using a Precellys homogenizer. Lysates were centrifuged for 10 min full speed at 4°C and transferred to a new tube. Protein concentration in the supernatant was determined using a BCA Protein Assay Kit (Thermo Scientific, #23225) and 500 μg of protein were used for the RBD assay. The following steps were performed using the Cytoskeleton Ras-activation assay biochem kit (Cytoskeleton, #BK008) according to the manufacturer’s instructions. Briefly, 30 ul of RAF-RBD beads were added to each tube and incubated to 1 h on a rotator at 4 °C. RAF-RBD beads were centrifuged for 1 min at 5000 g and 4 °C. The supernatant was removed, and beads were washed with 500 ul Wash Buffer and centrifuged for 3 min at 5000 g and 4 °C. Supernatant was carefully removed and 20 ul of 2 x Laemmli sample buffer was added to each tube. Samples were boiled to 5 min at 95 °C. Samples were loaded onto 4-12 % gradient pre-cast gel (Novex) and processed as Western blotting samples as described above. GDP incubated samples served as negative control, GTPγS served as positive control and His-tagged RAS protein was served as control protein. Total protein input served as loading control.

### mRNA isolation, qRT-PCR and RNA-Sequencing

PDAC tissue samples preserved in RNAlater (Sigma Aldrich #R0901) were homogenized with a Precellys homogenizer and RNA was extracted from tumour tissue using a QIAGEN RNAeasy kit (Quiagen #74104) according to the manufacturer’s instructions including DNase digestion. 0.5 µg of total RNA was reverse transcribed using a High-Capacity cDNA Reverse Transcription Kit (Thermo Scientific, #4368814) according to the manufacturer’s instructions. qRT-PCR was performed in technical replicates using QuantiFast SYBR Green RT-PCR Kit (Qiagen, # 204156) according to the manufacturer’s instructions. The reaction mixture without a template was run in duplicate as a control. *Gapdh* was used to normalise for differences in RNA input. For RNA sequencing, RNA integrity was analysed using the NanoChip (Agilent RNA 6000 Nano kit, 5067-1511) and 1 µg of RNA was prepared in water to a final volume of 50 µl. RNA sequencing was performed by the CRUK Beatson Institute’s Molecular Technologies Services using an Illumina TruSeq RNA sample prep kit, then run on an Illumina NextSeq using the High Output 75 cycles kit (2 x 36 cycles, paired end reads, single index). The raw sequence quality was assessed using the FastQC algorithm version 0.11.8. Sequences were trimmed to remove adaptor sequences and low-quality base calls, defined by a Phred score of <20, using the Trim Galore tool version 0.6.4. The trimmed sequences were aligned to the mouse genome build GRCm38.98 using HISAT2 version 2.1.0, then raw counts per gene were determined using FeatureCounts version 1.6.4. Differential expression analysis was performed using the R package DESeq2 version 1.22.2, and principal component analysis was performed using R base functions.

### Statistical analyses

Statistical analyses were performed using Microsoft Excel 2016 or GraphPad Prism (version 9.1). For survival analysis statistical comparisons were performed using the Mantel-Cox (long-rank) test. Box plots depict the interquartile range, median is indicated by the central line, and whiskers indicate minimum and maximum values. Error bars and the relevant statistical test are indicated in each figure legend.

## Results

### Loss of wild-type *Kras* facilitates pancreatic tumour initiation by oncogenic KRAS

In patients with pancreatic cancer (APGI group previously published in Bailey et al.^20^) the occurrence of allelic imbalance at the *KRAS* locus is associated with reduced survival (Figure 1A) and early progression as previously reported ^2,3^. In pancreatic cancer the proportion of patients with *KRAS* allelic imbalance is 40.5% (Figure 1B.) Previous work from the Rad group had shown high propensity for allelic imbalances of *KRAS* in human PDAC, and strong indication for such imbalances in pancreatic intraepithelial neoplasia (henceforth referred to as PanIN) progression ^21^.To better understand this process *in vivo*, we assessed the impact of *Kras* LOH upon both development of benign, pancreatic precursor lesions, known as PanIN, and ultimately development of pancreatic cancer.

**Figure 1:**
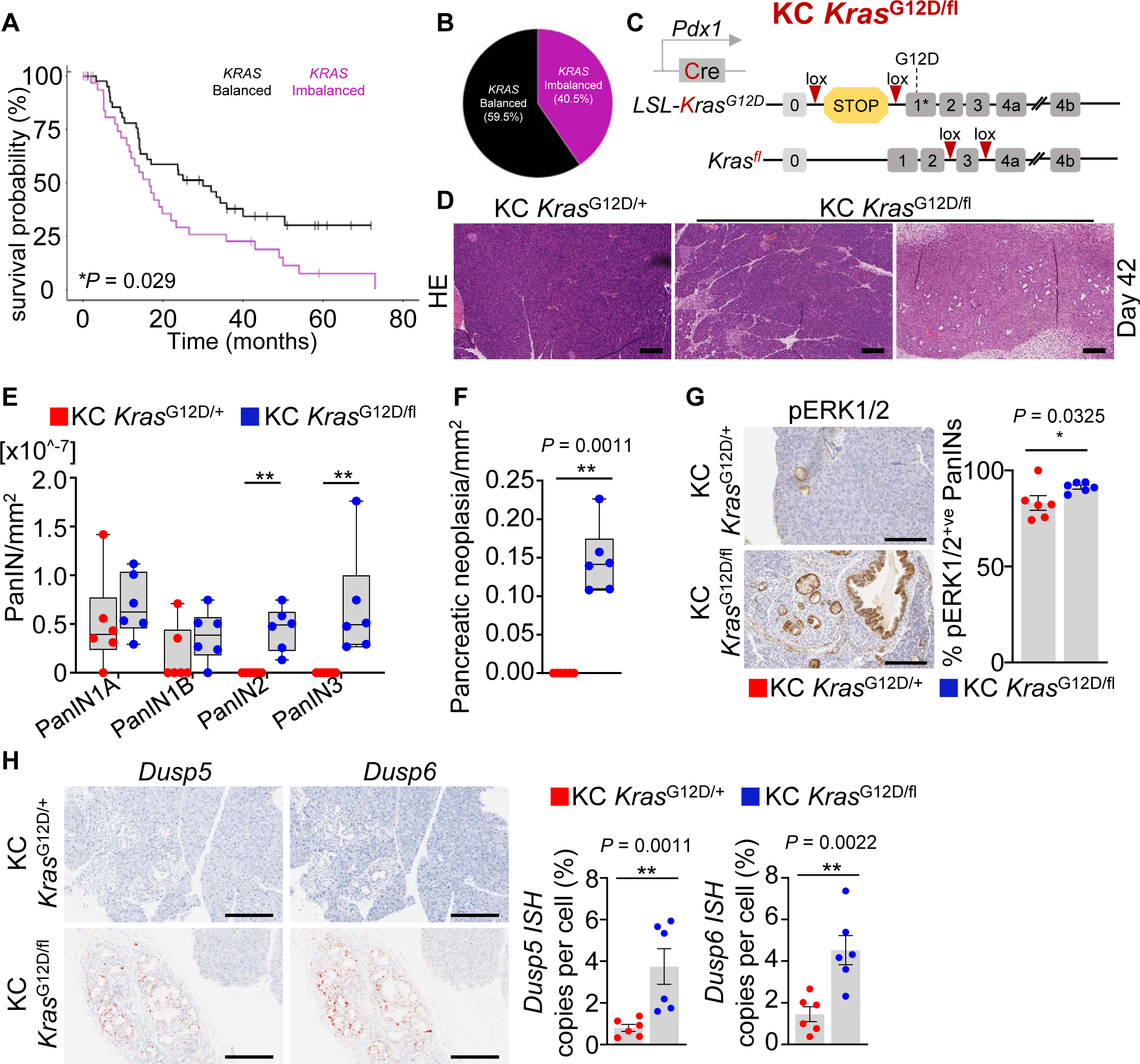
Loss of wild-type *Kras* increases PanIN formation in the presence of oncogenic *Kras*. a) Kaplan–Meier survival curve for human PDAC patients with *KRAS* alleles balanced and *KRAS* alleles imbalanced. KRAS balanced (n = 47), KRAS imbalanced (n = 32). \**P* = 0.029, log-rank (Mantel–Cox) test. b) Proportion of human pancreatic cancer patients with *KRAS* alleles balanced (59.5%) and *KRAS* alleles imbalanced (40.5%). c) Schematic representing generation of KC *Kras*^G12D/fl^ mice. Cre: Cre recombinase; loxP: Cre-loxP recombination site; KC: *Kras*^G12D/+^, Pdx1-Cre. d) Representative haematoxylin and eosin (H&E) images of pancreata from KC *Kras*^G12D/+^ and KC *Kras*^G12D/fl^ mice at 42 days of age. Scale bar 200 µm. e) Quantification of pancreatic intraepithelial neoplasia (PanINs) from KC *Kras*^G12D/+^ and KC *Kras*^G12D/fl^ mice at 42 days of age from whole haematoxylin and eosin (H&E) sections (n = 6 per group). Boxes depict interquartile range, central line indicates median and whiskers indicate minimum/maximum values. \*\**P* <0.01, one-way Mann–Whitney U test. f) Quantification of the area of pancreatic neoplasia per mm^2^ pancreas over one whole H&E section from KC *Kras*^G12D/+^ and KC *Kras*^G12D/fl^ mice at 42 days of age (*n* = 6 per group). Boxes depict interquartile range, central line indicates median and whiskers indicate minimum/maximum values. \*\**P* <0.0011, one-way Mann–Whitney U test. g) Left: Representative immunohistochemistry images of pERK1/2 of pancreata from KC *Kras*^G12D/+^ and KC *Kras*^G12D/fl^ mice at 42 days of age. Representative of six mice per group. Scale bar 200 µm. Right: Bar graphs showing quantification of pERK1/2 positive cells of the pancreatic epithelium of KC *Kras*^G12D/+^ and KC *Kras*^G12D/fl^ mice sampled at day 42 (KC *Kras*^G12D/+^, n = 6; KC *Kras*^G12D/fl^, n = 6). Data are mean ± s.e.m, \**P* = 0.0325, one-way Mann–Whitney U test. h) Left: Representative images of *in situ* hybridisation (*ISH*) of *Dusp5* and *Dusp6* of pancreata from KC *Kras*^G12D/+^ and KC *Kras*^G12D/fl^ mice at 42 days of age. Representative of six mice per group. Scale bar 200 µm. Right: Quantification of *Dusp5* and *Dusp6 ISH* staining of pancreatic lesions (PanINs) of pancreata from KC *Kras*^G12D/+^ and KC *Kras*^G12D/fl^ mice at 42 days of age (KC *Kras*^G12D/+^, n = 6; KC *Kras*^G12D/fl^, n = 6). Data are mean ± s.e.m, \*\**P* = 0.0011 (*Dusp5*), \*\**P* = 0.0022 (*Dusp6*), one-way Mann–Whitney U test.

To this end, we modelled precancerous pancreatic lesions, and development of pancreatic cancer by targeting the mouse developing pancreas with a conditional oncogenic LSL-*Kras*^G12D^ allele regulated by a constitutively expressed Cre-recombinase, under the control of the *Pdx1* promoter, commonly referred to as *Pdx1*-Cre *Kras*^G12D/+^ (henceforth referred to as KC *Kras*^G12D/+^) ^14^. Histologically, PanIN formation in the pancreata of these mice can be readily observed from around 10 weeks of age, with low penetrance of PDAC development observed at longer latencies of 12-16 months ^13,22^. Given that this process is driven by mutant KRAS, we initially sought to determine whether it could be modified by deletion of the wild-type *Kras* counterpart. To model this, we interbred *Pdx1*-Cre *Kras*^G12D/+^ mice with mice harbouring the conditional *Kras*^fl^ allele, thus generating *Pdx1*-Cre *Kras*^G12D/fl^ mice (henceforth referred to as KC *Kras*^G12D/fl^) (Figure 1C).

First, we performed a histological quantification of the number and stage of precancerous PanIN found in the pancreata of KC *Kras*^G12D/+^ and KC *Kras*^G12D/fl^ mice at 42 days of age. Strikingly, an increase in both number and grade of PanINs, along with extensive pancreatic neoplasia was observed in the pancreata of KC *Kras*^G12D/fl^ mice when compared to KC *Kras*^G12D/+^ mice (Figure 1D, E and F). Here, we defined pancreatic neoplasia, observed in KC *Kras*^G12D/fl^ mice, as replacement of the pancreatic parenchyma with a PanIN burden, along with immune and stromal infiltration. Pancreatic neoplasia was scored by area, whereas isolated PanIN lesions were scored individually by stage.

It is widely known that expression of oncogenic KRAS activates cellular growth arrest through downstream signalling of tumour suppressor pathways ^23^. An equivalent nuclear accumulation of p53 and p21 was observed in the PanIN epithelium of both pancreata of KC *Kras*^G12D/fl^ and KC *Kras*^G12D/+^ mice sampled at the 42-day timepoint (Supplementary Fig 1A). Moreover, few Ki67 positive cells were observed in the epithelium of PanINs in either model. It was notable that more Ki67 positive cells were detected within the pancreata of KC *Kras*^G12D/fl^ mice when compared to KC *Kras*^G12D/+^, however these were found within the stromal and inflammatory component of the lesions (Supplementary Fig 1A). This suggests that the substantial increase in early lesion formation in the pancreata of KC *Kras*^G12D/fl^ mice is not due to escape from long-term growth arrest or senescence.

Activated, oncogenic KRAS signalling in PDAC development is directly associated with activation of the downstream mitogen activated protein kinase (MAPK) pathway ^24^. PanINs in both KC *Kras*^G12D/fl^ and KC *Kras*^G12D/+^ models were positive for pERK1/2, indicative of activated MAPK signalling (Figure 1G). In addition, we focused on analysing the expression of two well-characterized MAPK dependent transcripts by *in situ* hybridisation (RNA*ish*) against *Dusp5* and *Dusp6* on pancreata from KC *Kras*^G12D/fl^ and KC *Kras*^G12D/+^ mice sampled at 42 days, showed higher expression of both *Dusp5* and *Dusp6* in PanINs from KC *Kras*^G12D/fl^ mice when compared to KC *Kras*^G12D/+^ controls (Figure 1H), suggesting that loss of the wild-type *Kras* in our model induces MAPK signalling, and drives a transcriptional response in the developing lesion.

Accumulation of PanINs, ADM and pancreatic neoplasia was observed in KC *Kras*^G12D/fl^ mice, accompanied by a fibroinflammatory reaction composed of stromal and immune infiltrate, observed by stroma development surrounding the architecture of advanced PanIN lesions with fibroinflammatory response shown by picrosirius red and Tenascin C positivity, markers for collagen/extracellular matrix deposition ^25,26^ (Supplementary Figure 1B). In addition, an increased abundance of α-smooth muscle actin (αSMA) a marker for activated fibroblasts was observed surrounding the epithelium of PanINs in KC *Kras*^G12D/fl^ compared to KC *Kras*^G12D/+^ mice (Supplementary Figure 1B). An abundant mucin content in the cytoplasm of PanINs of KC *Kras*^G12D/fl^ and KC *Kras*^G12D/+^ mice was demonstrated by Alcian Blue/Periodic acid-Schiff (AB/Pas) a mucin-specific stain.

### Deletion of wild-type *Kras* sensitizes pancreatic precursor lesions to MEK1/2 inhibition in early treatment intervention

Given the drastically increased PanIN burden and MAPK signalling in the pancreata of KC *Kras*^G12D/fl^ mice, we next sought to determine whether these PanIN lesions were sensitive to therapeutic targeting of the MAPK pathway. To this end, an early intervention study was performed using a clinically relevant inhibitor of the MAPK pathway intermediate kinases, MEK1/2 (AZD6244/selumetinib). As KC *Kras*^G12D/fl^ mice developed substantial PanIN burden by 42 days (Figure 1D, E & F), mice were treated with AZD6244 (25mgkg^-1^, twice daily by oral gavage) from 42 days of age, and sampled at a 70 day timepoint (Figure 2A). Strikingly, the pancreata of these AZD6244 treated KC *Kras*^G12D/fl^ mice exhibited an apparent reversion of the previously described PanIN initiation, with profoundly reduced area of pancreatic neoplasia when compared to vehicle treated mice and robust loss of pERK1/2 levels and reduced *Dusp6* expression (Figure 2B, C and D). No change was observed in proliferation in PanINs following AZD6244 treatment compared to control (Figure 2B). KC *Kras*^G12D/fl^ mice had high MYC levels and after treatment with MEK1/2 inhibitor cMYC levels were reduced (Figure 2D).

**Figure 2:**
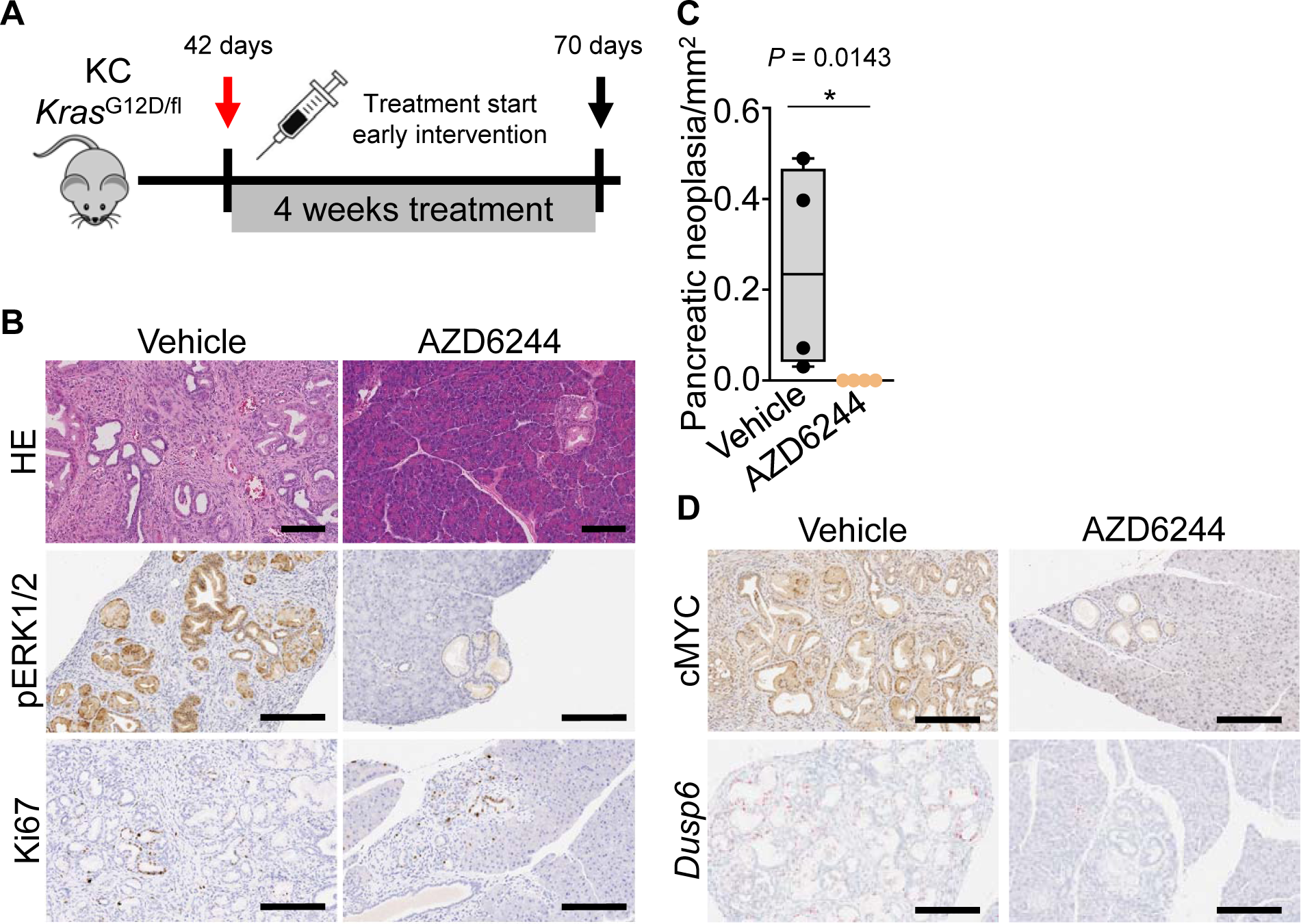
Early intervention with AZD6244 reduces PanIN burden in *Kras*^G12D/fl^ mice. a) Experimental set up: KC *Kras*^G12D/fl^ were treated from day 42 for 28 days with vehicle or AZD6244 and sampled at day 70. b) Representative H&E, pERK1/2 and Ki67 immunohistochemistry images of KC *Kras*^G12D/fl^ mice at 70 days of age treated with vehicle or AZD6244 as indicated for 28 days. Representative of four mice per group. Scale bar 200 µm. c) Quantification of the area of pancreatic neoplasia per mm^2^ pancreas over one whole H&E section of KC *Kras*^G12D/fl^ mice treated as indicated from day 42 for 28 days (n = 4 per group). Boxes depict interquartile range, central line indicates median and whiskers indicate minimum/maximum values. \**P* = 0.0143, one-way Mann–Whitney U test. d) Representative images of c-MYC immunohistochemistry and *Dusp6 in situ* hybridisation (*ISH*) of KC *Kras*^G12D/fl^ mice treated with vehicle or AZD6244 as indicated from day 42 for 28 days. Representative of four mice per group. Scale bar 200 µm.

To understand whether the profound reduction in pancreatic neoplasia burden following MEK inhibition is sustained and if the presence of oncogenic KRAS is enough to facilitate PanIN progression after withdrawal of MEK inhibition, we performed the early intervention approach as described above and followed this with a treatment intervention (“break”) for 7 days. In this setting, mice were treated with vehicle or AZD6244 for 28 days and then received no treatment for a further 7 days prior to analysis (Supplementary Figure 2A). Interestingly, even this short break in MAPK suppression was sufficient to allow a robust induction of pancreatic neoplasia formation in KC *Kras*^G12D/fl^ mice following withdrawal of MEK inhibition and the extent of neoplasia formation was similar to vehicle-treated KC *Kras*^G12D/fl^ mice (Supplementary Figure 2B, C). These data support our hypothesis that allelic imbalance at the *Kras* locus with oncogenic *Kras*^G12D^ mutation, results in substantial initiation of early pancreatic lesions, primarily in a MAPK dependent manner.

Our data above suggest that loss of wild-type KRAS in the context of an oncogenic KRAS^G12D^ mutant can facilitate early tumour initiation, in contrast prior studies have demonstrated that in certain circumstances, wild-type KRAS can act as a tumour suppressor ^27^. Therefore, we questioned whether wild-type KRAS can act as a tumour suppressor and the loss of wild-type KRAS can drive progression to pancreatic cancer. To achieve this, we aged KC *Kras*^G12D/fl^ and KC *Kras*^G12D/+^ mice to a clinical endpoint associated with ill health (median 200 days). When sampled at clinical endpoint, KC *Kras*^G12D/fl^ mice did not present with PDAC, rather the majority were sampled due to poor body condition. While the pancreata of these mice appeared macroscopically normal, histologically, they were characterised by substantial PanIN burden and the presence of pancreatic neoplasia, albeit in the absence of invasive PDAC. Analysis of KC *Kras*^G12D/fl^ and KC *Kras*^G12D/+^ mice at different timepoints (day 28, day 42, day 160 and day 190) showed that KC *Kras*^G12D/fl^ mice were characterised by substantial development of PanIN and associated fibrosis, gradually taking over the healthy acinar tissue at a much higher rate than in control KC *Kras*^G12D/+^ mice (Supplementary Figure 2D). The relative lack of PDAC development in the KC *Kras*^G12D/fl^ mice, particularly in the context of dramatically increased PanIN burden indicates that loss of wild-type *Kras* facilitates initiation and early lesion formation in the presence of an oncogenic *Kras*^G12D^ mutation, but is not sufficient for tumour progression. Thus, in the pancreas wild-type *Kras* does not act as a conventional tumour suppressor.

### Loss of wild-type *Kras* promotes oncogenic *Kras*-driven pancreatic tumourigenesis via increased MAPK signalling

Having observed that loss of wild-type KRAS in the context of mutated *Kras*^G12D^ was not sufficient to drive PDAC in isolation, we next sought to determine whether additional mutations in other canonical tumour suppressors might allow escape from the observed growth arrest-like phenotype and drive more penetrant pancreatic cancer. We chose to mutate P53 as this was still high in PanINs of *Kras*^G12D/fl^ mice and mutation is known to allow senescence escape and drive tumour progression. Therefore, we interbred both KC *Kras*^G12D/fl^ and KC *Kras*^G12D/+^ mice with a mouse carrying an oncogenic mutant *Trp53*^R172H/+^ allele, which encodes a dominant negative mutant version of p53, equivalent to one commonly found in human pancreatic cancers, thus generating Pdx1-Cre *Kras*^G12D/fl^ *Trp53*^R172H/+^ mice (henceforth referred to as KPC *Kras*^G12D/fl^). In this experimental setting, KPC *Kras*^G12D/fl^ and the equivalent control mice, Pdx1-Cre *Kras*^G12D/+^ *Trp53*^R172H/+^ (henceforth referred to as KPC *Kras*^G12D/+^), were aged to clinical endpoint, defined by exhibition of clinical signs of PDAC such as abdominal swelling, jaundice, hunching, piloerection or loss of body conditioning. Given that a profound lesion initiation phenotype was observed in KC *Kras*^G12D/fl^ mice, tumour development was monitored in KPC *Kras*^G12D/fl^ and KPC *Kras*^G12D/+^ mice from 42 days of age using high-resolution ultrasound imaging (Figure 3A). Ultrasound detection and measurement of pancreatic tumour initiation and resulting tumour burden was performed on mice once weekly from 6 weeks of age until the experimental endpoint (Figure 3A). The earliest indication of pancreatic cancer formation in KPC *Kras*^G12D/fl^ mice was detected from 42 days of age, whereas in KPC *Kras*^G12D/+^ controls, tumour formation was detected at a median of 85 days, indicating that loss of wild-type *Kras* facilitates tumour initiation (Figure 3B, Supplementary Figure 3A). Despite the earlier tumour initiation, we found that KPC *Kras*^G12D/fl^ tumours grew at a comparably slower rate after initial detection of tumour formation (Figure 3B), however overall median pancreatic tumour free survival of KPC *Kras*^G12D/fl^ mice was significantly shorter (median of 112 days), compared to KPC *Kras*^G12D/+^ control mice (median of 136 days) (\**P* = 0.0396, Kaplan Meier) (Figure 3C). Importantly, KPC *Kras*^G12D/fl^ mice developed fewer metastases when compared to the control KPC *Kras*^G12D/+^ cohort (Figure 3D). In addition, the primary pancreatic tumours which arose in KPC *Kras*^G12D/fl^ mice appeared histopathologically undifferentiated, whereas those which arose in KPC *Kras*^G12D/+^ mice appeared more differentiated in nature (Supplementary Figure 3B).

**Figure 3:**
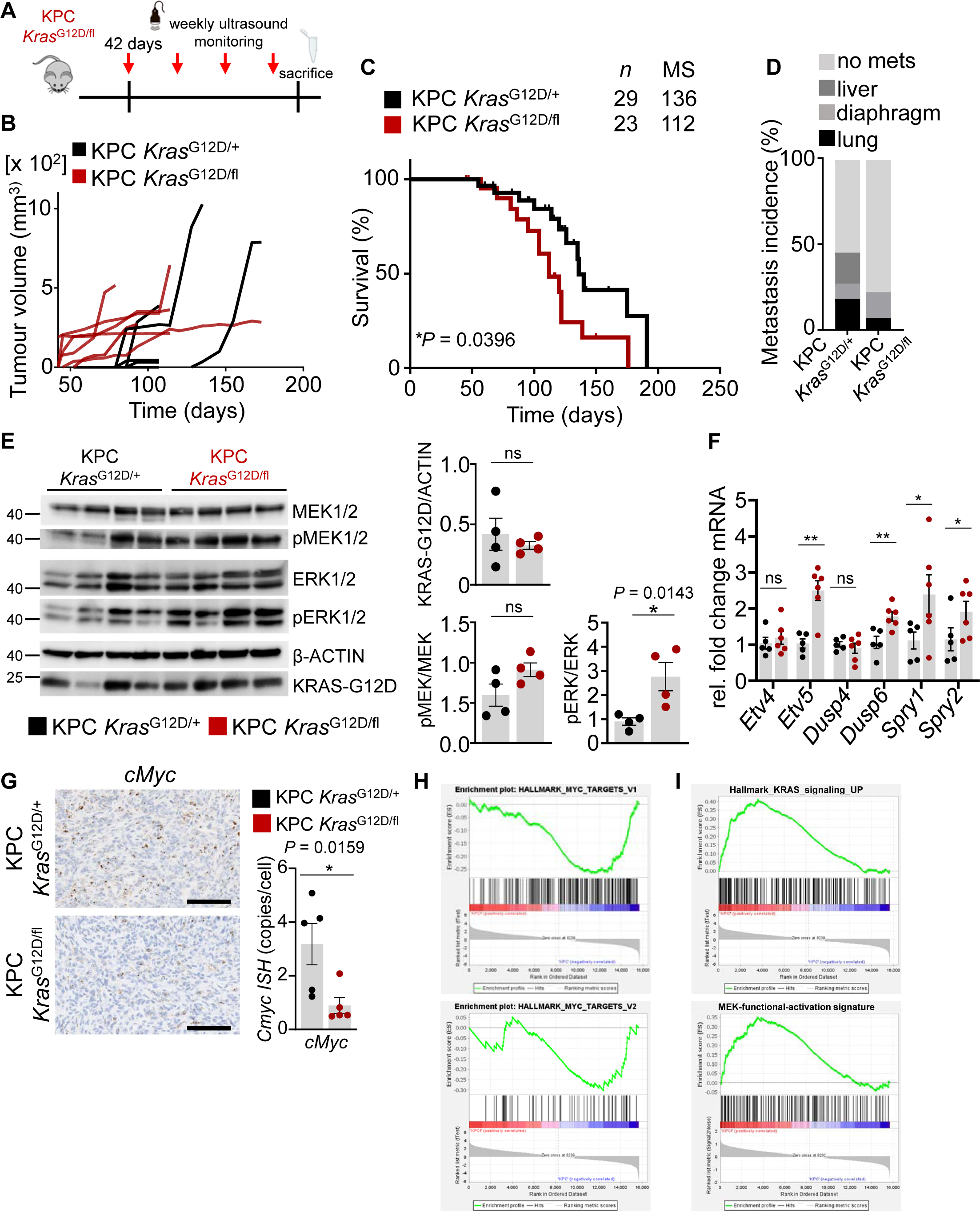
Loss of wild-type KRAS induces increased MAPK signalling in KPC *Kras*^G12D/fl^. a) Experimental schematic representing mice imaged once weekly by ultrasound from 6 weeks of age to clinical endpoint to follow tumour growth over time. b) Change of tumour volume (mm^3^) of KPC *Kras*^G12D/+^ and KPC *Kras*^G12D/fl^ mice aged to clinical endpoint measured once weekly by ultrasound imaging from 6 weeks of age. KPC *Kras*^G12D/fl^ (n = 5), KPC *Kras*^G12D/+^ (n = 5). c) Kaplan–Meier survival curves for KPC *Kras*^G12D/+^ and KPC *Kras*^G12D/fl^ mice aged to clinical endpoint. KPC *Kras*^G12D/fl^ (n = 29), KPC *Kras*^G12D/+^ (n = 23). \**P* = 0.0396 log-rank, (Mantel– Cox) test. Mice were censored when reaching clinical endpoint not associated with PDAC burden, such as primary tumours which arose from other tissue, mostly mucocutaneous papillomas or lymphomas (typically thymic), but also gastric and lung tumours. d) Incidence of metastasis (%) in KPC *Kras*^G12D/+^ and KPC *Kras*^G12D/fl^ mice. KPC *Kras*^G12D/fl^ (n = 27), KPC *Kras*^G12D/+^ (n = 20). e) Immunoblotting of pMEK1/2, MEK1/2, pERK1/2, ERK1/2, KrasG12D of KPC *Kras*^G12D/+^ and KPC *Kras*^G12D/fl^ PDAC tissue lysates generated from mice at clinical endpoint. ß-ACTIN was used as loading control. Each lane represents PDAC tissue from an individual mouse of the indicated genotype. Bar graph shows quantification of, pMEK1/2 levels, normalized to MEK1/2; pERK1/2 levels, normalized to ERK1/2; and KRAS-G12D levels normalized to β-ACTIN. Data represent mean ± s.e.m. \**P* = 0.0143, one-way Mann–Whitney U test. f) qRT-PCR analysis of *Etv4, Etv5, Dusp4, Dusp6, Spry1* and *Spry2* in KPC *Kras*^G12D/+^ and KPC *Kras*^G12D/fl^ PDAC tissue. Transcript levels were normalized to *Gapdh* (KPC *Kras*^G12D/+^ n = 5; KPC *Kras*^G12D/fl^ n = 6). Data represent mean ± s.e.m, *P* = 0.2 (*Etv4*), \*\**P* = 0.0087 (*Etv5*), *P* = 0.4 (*Dusp4*), \*\**P* = 0.0087 (*Dusp6*), \**P* = 0.0152 (*Spry1*), \**P* = 0.0411 (*Spry2*), one-way Mann–Whitney U test. g) Left: Representative *c-Myc in situ* hybridisation images of KPC *Kras*^G12D/+^ and KPC *Kras*^G12D/fl^ PDAC at clinical endpoint. Representative of 5 mice per group. Scale bar 200 µm. Right: Quantification of *c-Myc ISH* copies per cell in KPC *Kras*^G12D/+^ and KPC *Kras*^G12D/fl^ PDAC tissue (KPC *Kras*^G12D/+^ n = 5; KPC *Kras*^G12D/fl^ n = 5). Data represent mean ± s.e.m, \**P* = 0.0159, one-way Mann–Whitney U test. h) GSEA for Hallmark gene signatures for MYC targets genes V1 and V2 of PDAC tissue of KPC *Kras*^G12D/+^ and KPC *Kras*^G12D/fl^ mice at clinical endpoint. Y-axis is enrichment score (ES) and x-axis are gene represented in gene sets. i) GSEA for Hallmark gene signatures for “Kras signalling UP” and “MEK function activation” of PDAC tissue of KPC *Kras*^G12D/+^ and KPC *Kras*^G12D/fl^ mice at clinical endpoint. Y-axis is enrichment score (ES) and x-axis are gene represented in gene sets.

KPC *Kras*^G12D/fl^ tumours were not more proliferative than KPC *Kras*^G12D/+^, as indicated by BrdU incorporation in tumours (Supplementary Figure 3C and D). Accumulation of mutant p53 was reduced in KPC *Kras*^G12D/fl^ compared to KPC *Kras*^G12D/+^ tumours, correlating with the earlier shown lower incidence of metastasis in KPC *Kras*^G12D/fl^, as mutant p53 is required to drive a metastatic phenotype in murine PDAC^28^. In the more differentiated KPC *Kras*^G12D/fl^ tumours, the irregular tumour glands of PDAC appear to be surrounded by desmoplastic stroma, composed of stromal, inflammatory and immune cells along with the characteristic extracellular matrix. In contrast to the KC *Kras*^G12D/fl^ mice the stromal architecture appeared equivalent in KPC *Kras*^G12D/fl^ and KPC *Kras*^G12D/+^ tumours observed by αSMA and picrosirius red staining (Supplementary Figure 3C and D).

Given the apparent discordance between the early tumour initiation phenotypes, and subsequent tumour growth, we next sought to better understand the molecular impact that loss of wild-type *Kras* has in the context of an oncogenic *Kras*^G12D^ mutant in pancreatic tumours *in vivo*. First, we determined whether loss of wild-type *Kras* altered the proportion of GTP-bound (or “active”) KRAS found in KPC *Kras*^G12D/fl^ and KPC *Kras*^G12D/+^ tumours. Using RAF-RBD pulldowns we observed no changes in active KRAS, suggesting an equal proportion of active KRAS in KPC *Kras*^G12D/fl^ and KPC *Kras*^G12D/+^ tumours (Supplementary Figure 4A, B).

Despite the apparent lack of impact upon GTP binding, an upregulation of MAPK dependent transcripts was observed in early pancreatic lesions in the KC *Kras*^G12D/fl^ model. It was notable that these included transcripts encoding negative regulators of the MAPK pathway, comprising a well-established feedback mechanism. This observation is in line with a previously demonstrated link between oncogenic *KRAS* allelic imbalances, in particular LOH, and increased downstream signalling ^29^. Therefore, we investigated whether loss of the wild-type *Kras* allele in mouse pancreatic tumours can alter signalling through downstream effector pathways and activate any associated feedback loops. Tumours arising in KPC *Kras*^G12D/fl^ mice exhibited substantially increased pERK1/2 when compared to control KPC *Kras*^G12D/+^ tumours (Figure 3E). This pattern is in contrast to that seen in early initiating lesions, and suggests a robust upregulation of MAPK signalling in tumours arising in the KPC *Kras*^G12D/fl^ model *in vivo*, when compared to KPC *Kras*^G12D/+^ controls. Notably, other common effector pathways downstream of KRAS, such as the PI3K signalling pathway appeared unchanged (Supplementary Figure 4C). This induction of MAPK signalling in KPC *Kras*^G12D/fl^ tumours was supported by significantly increased expression of MAPK-dependent transcriptional targets such as *Etv5, Dusp6, Spry1* and *Spry2* in KPC *Kras*^G12D/fl^ tumours compared to controls (Figure 3F). In accordance, protein levels of DUSP6 were found to be elevated in KPC *Kras*^G12D/fl^ tumours compared to KPC *Kras*^G12D/+^ mouse tumours (Supplementary Figure 4C). These data suggest that in the context of a pancreatic tumour, loss of wild-type *Kras* potentiates oncogenic KRAS-driven MAPK signalling and transcription of downstream target genes. Interestingly, we observed a significant downregulation of *Cmyc ISH* levels following loss of wild-type *Kras* in KPC *Kras*^G12D/fl^ tumours compared to KPC *Kras*^G12D/+^ tumours (Figure 3G). This was associated with negative enrichment for MYC target genes V1 and V2 (Figure 3H). While PanIN lesions in KC *Kras*^G12D/fl^ exhibit high levels of cMYC at the protein level (Figure 2E), transcriptional expression of *Myc* and its target genes is reduced in tumours from KPC *Kras*^G12D/fl^ mice when compared to KPC *Kras*^G12D/+^ controls. These observations might suggest a selective pressure favouring MYC-low tumour cell populations during tumour progression, specifically in the context of elevated MAPK signalling ^30^.

Several recent studies have highlighted that transcriptional profiling can be a useful tool in understanding and the classification of human PDAC. Given the clear phenotypic differences between the KPC *Kras*^G12D/fl^ and KPC *Kras*^G12D/+^ models (Figure 3), we sought to determine whether transcriptional profiling of these tumours could be used to stratify and align to previously described human PDAC subtype classifications. Analysis of KPC *Kras*^G12D/fl^ and KPC *Kras*^G12D/+^ tumours using the Bailey classification suggested no significant enrichment for any subtype in both KPC *Kras*^G12D/+^ and KPC *Kras*^G12D/fl^ tumours (Supplementary Figure 4D) ^31,32^. However, gene programmes GP6, GP7, and GP8 from the Bailey classification, all of which are associated with immune cell specific gene expression signatures, were significantly enriched in tumours arising in KPC *Kras*^G12D/fl^ mice (Supplementary Figure 4E). These immune gene programme signatures include those representative of B-cell signalling pathways, antigen presentation, CD4^+^ and CD8^+^ T cell signalling pathways. Moreover, immune regulatory hallmark gene sets such as “interferon alpha response” and “interferon gamma response” were also significantly enriched in KPC *Kras*^G12D/fl^ tumours compared to KPC *Kras*^G12D/+^ (Supplementary Figure 4F). In addition to subtyping analysis, geneset enrichment analysis (GSEA) demonstrated that genes up-regulated by KRAS activation (Hallmark gene set “KRAS-signalling up”) and genes correlating with functional MEK activation (“Mean MEK-functional-activation signature gene expression”) were specifically enriched in KPC *Kras*^G12D/fl^ tumours ^33^ (Figure 3I).

### Loss of wild-type *Kras* sensitizes late-stage pancreatic tumours to MEK1/2 inhibition

Given that MAPK signalling was enriched in KPC *Kras*^G12D/fl^ tumours, we next investigated whether MEK inhibition is effective in the context of an established pancreatic tumour. This is of particular importance as the majority of pancreatic cancer patients present with late-stage, established and difficult to treat disease. To achieve this, cohorts of KPC *Kras*^G12D/fl^ and KPC *Kras*^G12D/+^ mice with palpable tumours confirmed by ultrasound, were enrolled into treatment groups and received AZD6244 or the vehicle control. Tumour growth was monitored once weekly by ultrasound imaging over the course of treatment (Figure 4A). Consistent with previous reports, we observed that control KPC *Kras*^G12D/+^ tumours were resistant to MEK1/2 inhibition. The AZD6244 or vehicle treated KPC *Kras*^G12D/+^ mice showed no overall difference in survival and both had a continuous, rapid and comparable tumour growth ^34^ (Figure 4B). Strikingly, ultrasound monitoring, and measurement of tumours demonstrated significant tumour shrinkage in KPC *Kras*^G12D/fl^ mice following enrolment on AZD6244 treatment, but not following vehicle treatment (Figure 4B). This tumour shrinkage was detectable within the first 7 days of treatment and the tumour volume typically remained stable or further shrank for up to 21 days after beginning treatment. After this 21-day period, therapeutic resistance was acquired, tumours progressed, and grew rapidly (Figure 4C). A profound extension of survival was observed in KPC *Kras*^G12D/fl^ mice treated with AZD6244 compared to vehicle treated control mice (Figure 4D).

**Figure 4:**
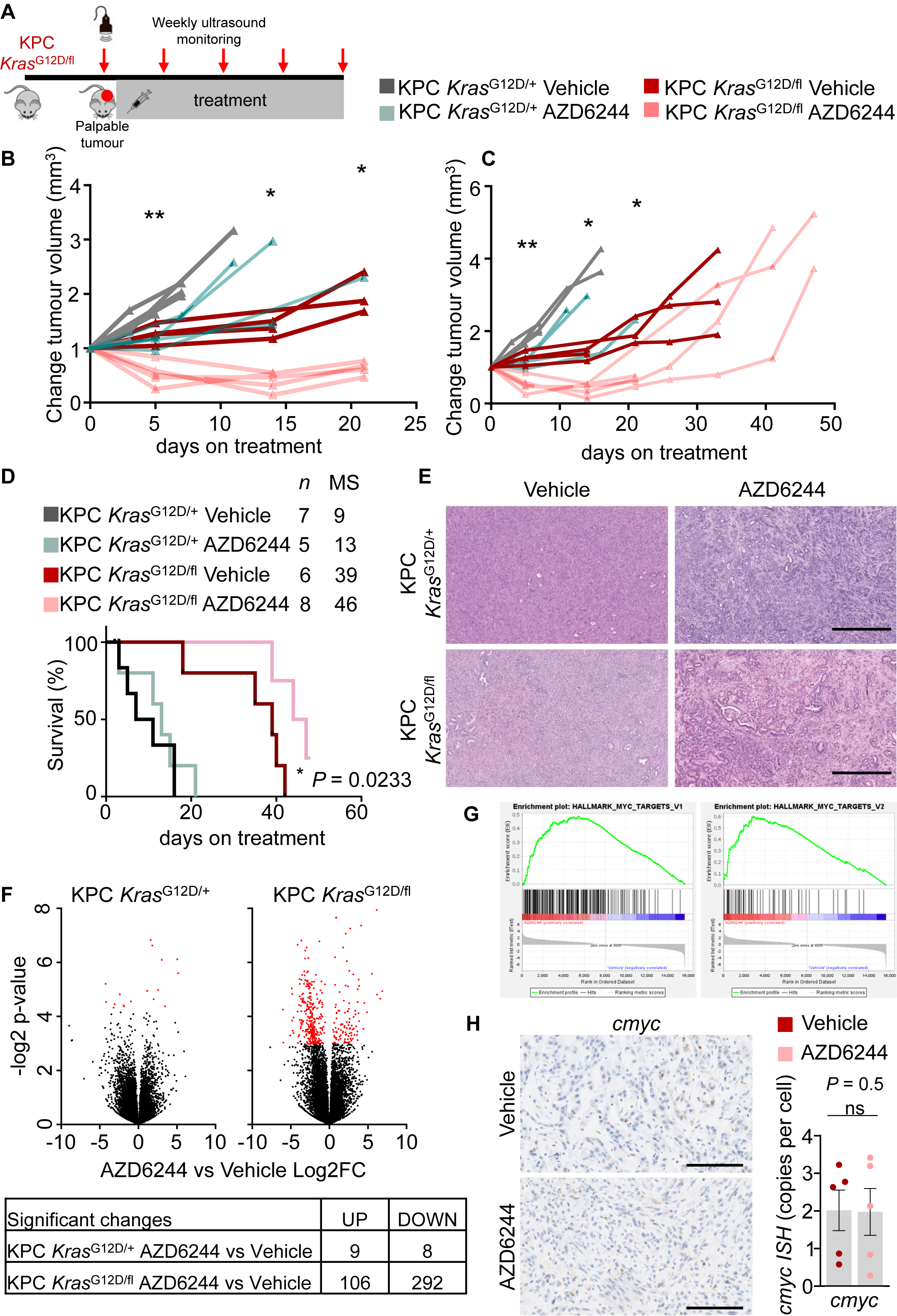
Late-stage intervention with MEK1/2 inhibition improves survival of KPC *Kras*^G12D/fl^ mice. a) Experimental set up: KPC *Kras*^G12D/+^ and KPC *Kras*^G12D/fl^ mice were palpated for tumour burden. Tumour burden in KPC *Kras*^G12D/+^ and KPC *Kras*^G12D/fl^ mice was confirmed by ultrasound imaging, mice were treated continuously from the following day with either vehicle or AZD6244, and tumour growth was monitored by ultrasound imaging once weekly to clinical endpoint. b) Change of tumour volume (mm^3^) of KPC *Kras*^G12D/+^ and KPC *Kras*^G12D/fl^ mice treated with either vehicle control or AZD6244 from palpable tumour and aged to clinical endpoint. Tumour volume was measured once weekly by high-resolution ultrasound imaging. Graph shows tumour volume for 3 weeks of treatment. Each line represents an individual mouse of the indicated genotype and treatment. \*\**P* = 0.079 (d7), \**P* = 0.0179 (d14), \**P* = 0.0286 (d21) KPC *Kras*^G12D/fl^ vehicle vs AZD6244. c) Change of tumour volume (mm^3^) of KPC *Kras*^G12D/+^ and KPC *Kras*^G12D/fl^ mice treated with either vehicle control or AZD6244 from palpable tumour and aged to clinical endpoint. Tumour volume was measured once weekly by ultrasound imaging. Graph shows tumour volume for full treatment cohort (from b). Each line represents an individual mouse of the indicated genotype and treatment. \*\**P* = 0.079 (d7), \**P* = 0.0179 (d14), \**P* = 0.0286 (d21) KPC *Kras*^G12D/fl^ vehicle vs AZD6244. d) Kaplan–Meier survival curves for KPC *Kras*^G12D/+^ and KPC *Kras*^G12D/fl^ mice treated with vehicle or AZD6244 as indicated from palpable tumour burden and aged to clinical endpoint. KPC *Kras*^G12D/fl^ vehicle (n = 6), KPC *Kras*^G12D/fl^ AZD6244 (n = 8), KPC *Kras*^G12D/+^ vehicle (n = 7), KPC *Kras*^G12D/+^ AZD6244 (n = 5). \**P* = 0.0233 log-rank (Mantel–Cox) test. e) Representative H&E images of for KPC *Kras*^G12D/+^ and KPC *Kras*^G12D/fl^ mice treated with vehicle or AZD6244 as indicated from palpable tumour burden and aged until clinical endpoint. Representative of five mice per group. Scale bar 500µm. f) Left: Volcano blot for differentially expressed genes of KPC *Kras*^G12D/+^ tumours treated with vehicle or AZD6244 from palpable tumour burden to clinical endpoint. Significantly altered genes are marked in red. Right: Volcano blot for differentially expressed genes of KPC *Kras*^G12D/fl^ tumours treated with vehicle or AZD6244 from palpable tumour burden to clinical endpoint. Significantly altered genes are marked in red. g) GSEA for Hallmark gene signatures for MYC targets genes V1 and V2 KPC *Kras*^G12D/+^ and KPC *Kras*^G12D/fl^ treated as indicated with either vehicle or AZD6244 from palpable tumour burden to clinical endpoint. Y-axis is enrichment score (ES) and x-axis are gene represented in gene sets. h) Left: Representative images of *c-Myc in situ* hybridisation of KPC *Kras*^G12D/fl^ mice treated with either Vehicle or AZD6244 from palpable tumour burden as indicated to clinical endpoint. Scale bar 200 µm. Right: Quantification of *c-Myc ISH* copies per cell in KPC *Kras*^G12D/fl^ treated with either Vehicle or AZD6244 from palpable tumour burden to clinical endpoint (*Kras*^G12D/fl^ Vehicle n = 5; KPC *Kras*^G12D/fl^ AZD6244 n = 5). Data represent mean ± s.e.m, *P* = 0.5, one-way Mann–Whitney U test.

The therapeutic response in KPC *Kras*^G12D/fl^ tumours treated with AZD6244 was associated with histopathological changes when compared to vehicle treated ones (Figure 4E). While vehicle treated tumours appeared to exhibit the undifferentiated morphology observed in the parental KPC *Kras*^G12D/fl^ cohort, tumours from the AZD6244 treated group exhibited a more well-differentiated morphology more reminiscent of the KPC *Kras*^G12D/+^ cohort (or indeed, KPC *Kras*^G12D/+^ treated with AZD6244 or vehicle).

To gain further insight into mechanisms which drive sensitivity to AZD6244 in KPC *Kras*^G12D/fl^ tumours, and thus potentially identify targets or pathways which might sensitise the control KPC *Kras*^G12D/+^ tumours, we performed transcriptomic analysis of tumours from KPC *Kras*^G12D/fl^ or KPC *Kras*^G12D/+^ cohorts treated with either AZD6244 or vehicle from palpable tumour detection. In the AZD6244 treatment resistant KPC *Kras*^G12D/+^ tumours, AZD6244 treatment induced very few significantly differentially regulated genes groups (9 significantly upregulated and 8 significantly downregulated genes) (Figure 4F). In contrast, in KPC *Kras*^G12D/fl^ tumours, which were sensitive to AZD6244 treatment (MEK1/2 inhibition), there were a substantially higher number of differentially regulated genes in AZD6244 treated vs vehicle treated groups (106 significantly upregulated and 292 significantly downregulated genes) (Figure 4F). Transcriptional analysis for the “MEK-functional-activation” gene set revealed no transcriptional enrichment in KPC *Kras*^G12D/+^ cohorts treated with either AZD6244 or vehicle, supporting the observation of resistance to MEK inhibition in KPC *Kras*^G12D/+^. In KPC *Kras*^G12D/fl^ tumours treated with AZD6244, however, we observed enrichment for the “MEK-functional-activation” gene set, suggesting acquired resistance to MEK inhibition in KPC *Kras*^G12D/fl^ tumours at clinical endpoint (Supplementary Figure 5A) ^33^. Together, this transcriptional analysis reinforces observations made in KPC *Kras*^G12D/fl^ or KPC *Kras*^G12D/+^ mice treated with either AZD6244 or vehicle, supporting the observed resistance of KPC *Kras*^G12D/+^ tumours to MEK inhibition, by the small number of differentially expressed genes, and the initial response of KPC *Kras*^G12D/fl^ tumours to MEK inhibition with acquired resistance over time, by enrichment of activated MAPK signalling gene sets.

Transcriptional subtype classification of KPC *Kras*^G12D/+^or KPC *Kras*^G12D/fl^ derived tumours indicated that treatment with AZD6244 had no impact upon subtype in KPC *Kras*^G12D/+^ tumours, further highlighting the robust resistance to MEK inhibition in this setting, while, enrichment for the immunogenic subtype in KPC *Kras*^G12D/fl^ tumours was reversed by treatment with AZD6244 (Supplementary Figure 6A). This was associated with an apparent loss of enrichment of the immune signalling associated gene programmes GP6, GP7 and GP8 following AZD6244 treatment in this model (Supplementary Figure 6B) and negative enrichment for immune associated hallmark gene sets in KPC *Kras*^G12D/fl^ tumours treated with AZD6244 compared to vehicle treated counterparts (Supplementary Figure 5B).

In line with the apparent suppression of cMyc expression, and MYC target genes in KPC *Kras*^G12D/fl^ tumours, in these mice which have acquired resistance to AZD6244, and are subsequently sampled at clinical endpoint, we observe a significant positive enrichment for cMYC target gene expression (Figure 4G). Nonetheless, *Cmyc ISH* levels were not significantly changed at clinical endpoint in pancreata of KPC *Kras*^G12D/fl^ mice treated with either Vehicle or AZD6244 (Figure 4H). These observations might support the hypothesis that high levels of cMyc target gene expression in the context of elevated MAPK signalling might be deleterious to tumour growth, with suppression of MAPK signalling through MEK1/2 inhibition allowing outgrowth of tumour cells exhibited elevated cMyc target gene expression. The robust therapeutic response elicited by AZD6244 suggests that KPC *Kras*^G12D/fl^ tumours have increased dependency upon MAPK signalling and suggests that a therapeutic opportunity may exist in patients with allelic imbalance at the *KRAS* locus for MAPK targeting therapeutic approaches. The differential response to AZD6244 observed in KPC *Kras*^G12D/fl^ and KPC *Kras*^G12D/+^ tumours further supports the hypothesis that loss of the wild-type *Kras* allele profoundly affects the oncogenic signalling within pancreatic tumours, and that the presence of a wild-type *Kras* allele can contribute to therapeutic resistance in *Kras* mutant tumours.

## Discussion

For many years, research has been focused on understanding oncogenic driver genes and their functional impact in cancer. Recent studies provide evidence that not only the oncogenic *RAS* mutations but also the relative oncogene gene dosage due to allelic imbalance and copy number variations are modulators of oncogenic signalling output, tumour development and incidence of metastasis ^2,10,11^. Notably, there has been growing evidence that *RAS* relative gene dosage modulates response to therapy ^11^.

Our data provide evidence that loss of wild-type *Kras* plays a crucial role in PDAC biology and phenotypic diversification. We developed a genetically engineered mouse model of *Kras* loss of heterozygosity in pancreatic cancer. This study shows that early pancreatic cancer development was facilitated in wild-type *Kras* depleted mice in the context of oncogenic *Kras*, observed by increased numbers and more advanced precursor lesions (PanIN) and extensive pancreatic neoplasia in the pancreas of wild-type *Kras* deficient *Kras* mutant mice. Loss of wild-type *KRAS* in the presence of mutant *KRAS*^G12D^ has been shown to induce MAPK signalling as observed by transcriptional activation of MAPK regulators. We show that depletion of wild-type *Kras* in *Kras* mutant mice facilitates tumourigenesis in the pancreas yet was not sufficient to drive progression to invasive PDAC, suggesting that wild-type KRAS acts to restrain mutant KRAS signalling but does not act as tumour suppressor in pancreatic cancer.

Previous studies suggest that the presence of wild-type RAS modulates response to treatment in oncogenic RAS driven tumours ^11,29^. While wild-type KRAS proficient established tumours were resistant to MAPK inhibition with a MEK1/2 inhibitor, we observed a robust initial response to treatment in KPC *Kras*^G12D/fl^ tumours. Initial tumour shrinkage in KPC *Kras*^G12D/fl^ mice treated with AZD6244 is an important point to highlight, as many patients present clinically with advanced pancreatic cancers that is not suitable for surgical resection, which is the only effective cure. Therefore, even transient therapeutic response that drives tumour shrinkage in a defined subset of patients (patients with *KRAS* LOH) might offer a promising treatment opportunity and potential clinical benefit in the neoadjuvant setting. Similar approaches are currently in phase-II clinical trials with neoadjuvant chemotherapy prior to surgical resection in pancreatic cancer (PRIMUS-002) ^35^.

Wild-type *Kras* deletion had profound effects on the transcriptional landscape of tumours, suggesting phenotypic alteration in tumours deficient for wild-type *Kras*. A specific enrichment for the immunogenic subtype was observed in the KPC *Kras*^G12D/fl^ tumours, along with upregulation of immune cell specific gene programmes. Given the enrichment for immunogenic gene programmes, this may suggest that specific a subset of patients with LOH *Kras* allelic imbalance may benefit from immune checkpoint inhibitors such as anti-PD1/PD-L1 or anti-CTLA4 targeting therapies, which should be the subject of further investigation.

In summary, these data demonstrate that even simple allelic imbalances at the *Kras* locus can have a profound impact upon all aspects of the tumourigenic process, encompassing initiation, progression and therapeutic sensitivity. These data in turn suggest that the presence of a wild-type copy of *Kras* can act to suppress the impact of oncogenic *Kras*. We have gone on to show that preclinical models featuring allelic imbalance at the *Kras* locus in the context of oncogenic *Kras* may represent a subset of pancreatic cancer patients in the clinic. Finally, we demonstrate significant therapeutic efficacy of clinically relevant inhibitors of the MAPK pathway in this preclinical setting, hinting at the possibility of clinical efficacy in a substantial subset of patients.

Altogether, the data presented in this study highlight that mutant and wild-type *Kras* allelic status in *Kras*^G12D^-driven pancreatic cancer has important implications on PDAC biology and therapeutic response. Poor response of patients to therapeutic intervention may be explained by mutant and wild-type *Kras* allelic status. We demonstrate that stratification of patients by *Kras* allelic imbalances (loss of heterozygosity, and mutant *Kras* copy number) might aid to identify therapeutic vulnerabilities and lead to more patient specific treatment modalities and overall contribute to improved treatments responses and patient care.

## Author contributions

S.K.F, A.K.N., A.D.C and O.J.S. designed the research. S.K.F., A.K.N. and C.F. performed experiments and analysed data. S.K.F., A.K.N., K.G. and W.C., generated and analysed transcriptomic data. S.K.F., A.K.N. and C.N. performed immunohistochemistry. R.U.-G. and D.C. provided patient data. S.B. provided AZD6244. J.P.M. provided expertise and provided feedback. S.K.F., A.K.N., A.D.C. and O.J.S. wrote the paper, all authors read the manuscript and provided critical comments.

## Acknowledgements

The authors are grateful to the CRUK Beatson Institute Central Services, Histology Services, Transgenics and BSU for technical support and all members of the Sansom lab for discussion of the data and the manuscript. O.J.S. was supported by CRUK grants A21139, A12481, A17196 and A31287 and ERC Starting grant 311301. A.K.N. was supported by a Novartis grant awarded to O.J.S. S.K.F. was supported by a Pancreatic Cancer UK Future Leaders Academy studentship - awarded to O.J.S and J.P.M. K.G. was supported by CRUK Grand Challenge Specificancer Grand Challenge Consortium (A29055 awarded to O.J.S.). C.F, C.N, W.C, and A.D.C. were supported by CRUK Beatson Institute core funding A17196, A31287 – awarded to O.J.S. J.P.M was supported by CRUK Beatson Institute core funding A17196, A31287 and A29996.

---

**Supplementary Figure 1:**
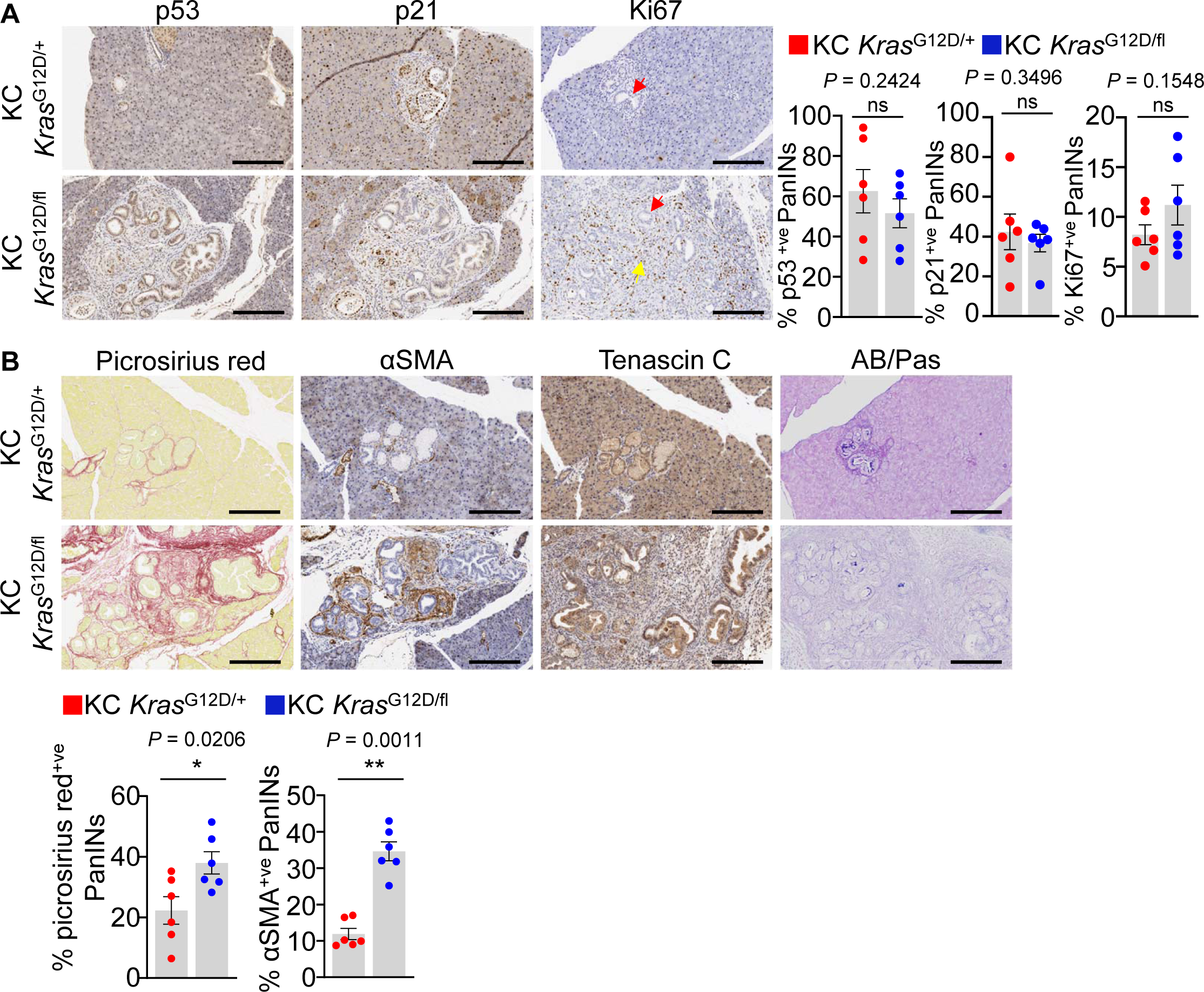
Increased fibrosis in wild-type *Kras* deficient PanINs. a) Left: Representative immunohistochemistry images of p53, p21 and Ki67 of pancreata from KC *Kras*^G12D/+^ and KC *Kras*^G12D/fl^ mice at 42 days of age. Red arrow indicates PanIN lesions (epithelial) and yellow arrow indicates Ki67 positive cells in the stromal compartment. Representative of six mice per group. Scale bar 200 µm. Right: Quantification of p53, p21 and Ki67 positive pancreatic lesions (PanINs) of pancreata from KC *Kras*^G12D/+^ and KC *Kras*^G12D/fl^ mice at 42 days of age (KC *Kras*^G12D/+^, n = 6; KC *Kras*^G12D/fl^, n = 6). Data are mean ± s.e.m, *P* = 0.2424 (p53), *P* = 0.3496 (p21), *P* = 0.1548 (Ki67), one-way Mann–Whitney U test. b) Top: Representative images of picrosirius and AB/PAs staining and immunohistochemistry of αSMA and Tenascin C of pancreata from KC *Kras*^G12D/+^ and KC *Kras*^G12D/fl^ mice at 42 days of age. Scale bar 200 µm. Bottom: Quantification of picrosirius red and αSMA positivity (%) of PanIN lesions of pancreata from KC *Kras*^G12D/+^ and KC *Kras*^G12D/fl^ mice at 42 days of age (KC *Kras*^G12D/+^, n = 6; KC *Kras*^G12D/fl^, n = 6). Data are mean ± s.e.m, \**P* = 0.0206 (picrosirius red), \*\**P* = 0.0011 (αSMA), one-way Mann–Whitney U test.

**Supplementary Figure 2:**
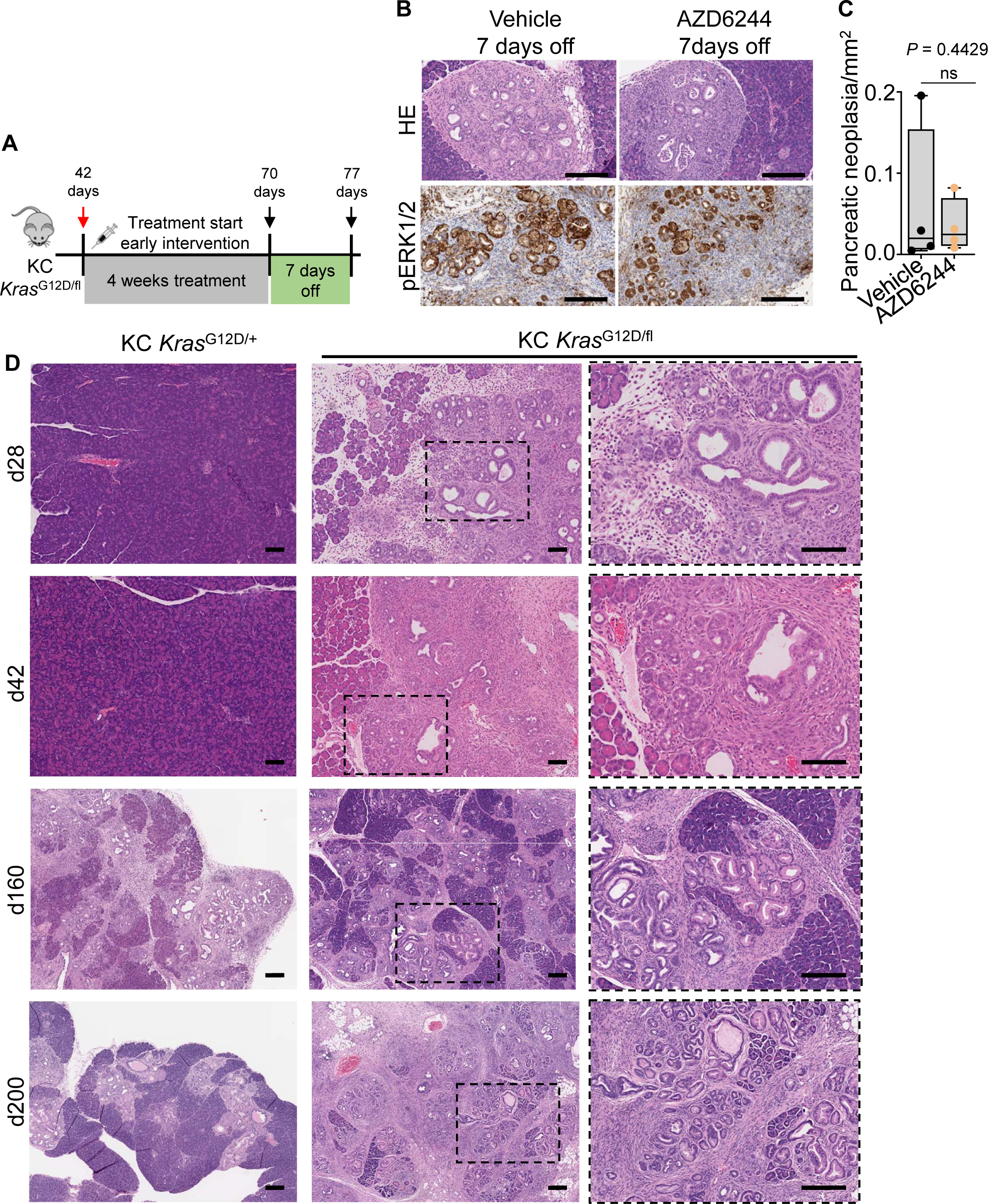
Retrieval of AZD6244 allows reformation of PanINs in KC *Kras*^G12D/fl^. a) Experimental set up: *Kras*^G12D/fl^ were treated from day 42 for 28 days with vehicle or AZD6244 as indicated prior to a 7-day drug holiday and sampling at day 77. b) Representative H&E and pERK1/2 images of KC *Kras*^G12D/fl^ mice at 77 days of age treated as indicated for 28 days and 7 days off. Representative of four mice per group. Scale bar 200 µm. c) Quantification of the area of pancreatic neoplasia per mm^2^ pancreas over one whole H&E section from KC *Kras*^G12D/fl^ mice at 77 days of age treated as indicated from day 42 for 28 days and 7 days off (n = 4 per group). Boxes depict interquartile range, central line indicates median and whiskers indicate minimum/maximum values. *P* = 0.4429, one-way Mann–Whitney U test. d) Representative H&E images of pancreata from KC *Kras*^G12D/+^ and KC *Kras*^G12D/fl^ mice of indicated timepoints. Dashed boxes show higher magnification. Scale bar 200 µm.

**Supplementary Figure 3:**
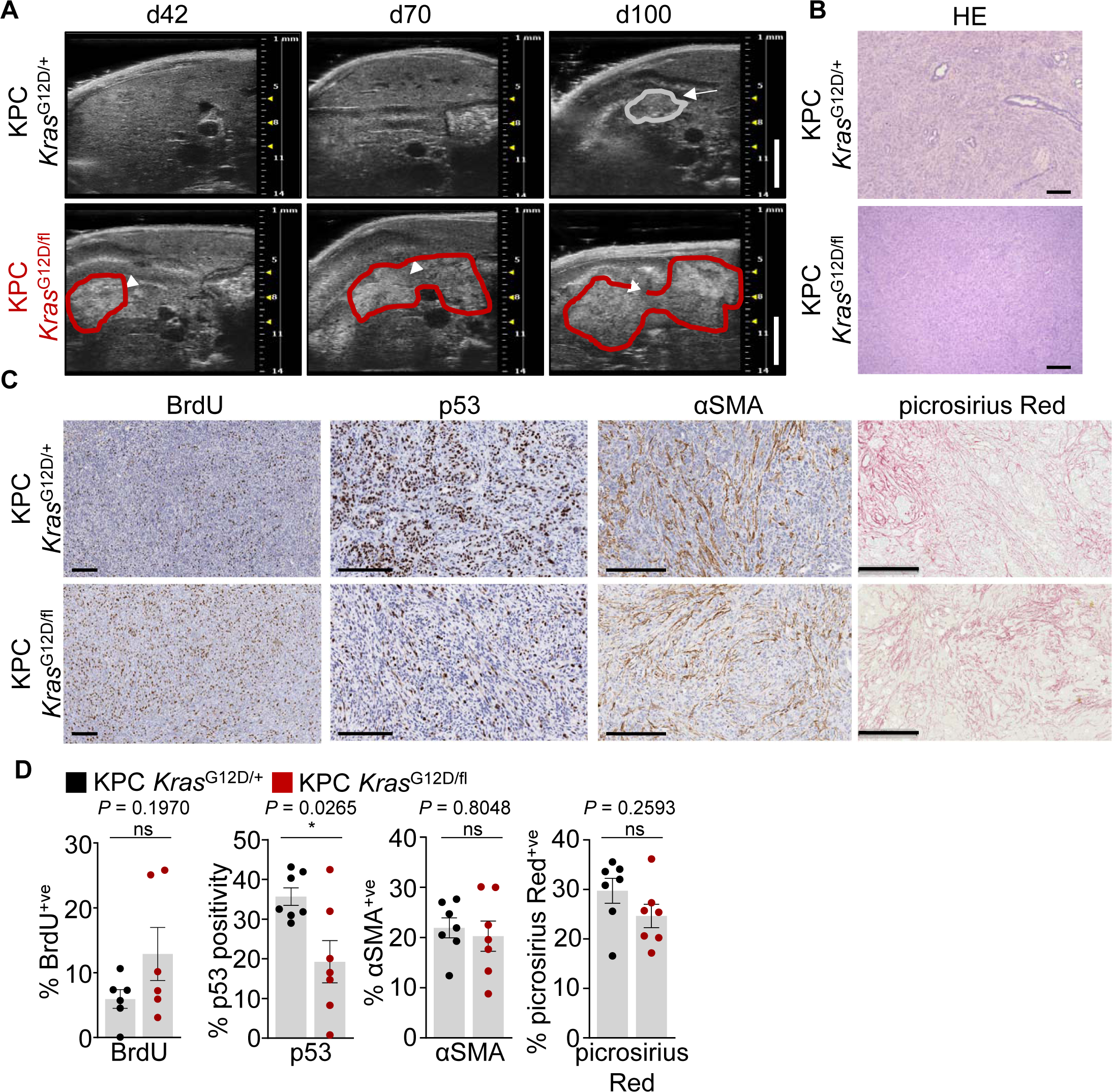
Acceleration of pancreatic tumour initiation after loss of wild-type *Kras* in KPC *Kras*^G12D/fl^ mice. a) Representative images of high-resolution ultrasound imaging of KPC *Kras*^G12D/+^ and KPC *Kras*^G12D/fl^ mice at day 42, day 70 and day 100. Arrows and mark up indicate PDAC. Representative of five mice per group. Scale bar 5 mm. b) Representative H&E images of KPC *Kras*^G12D/+^ and KPC *Kras*^G12D/fl^ mice aged to clinical endpoint. Scale bar 200 µm. c) Representative images of BrdU, p53, αSMA immunohistochemistry and picrosirius red staining of KPC *Kras*^G12D/+^ and KPC *Kras*^G12D/fl^ mice aged to clinical endpoint, (KPC *Kras*^G12D/+^, n = 7 (n = 6 for BrdU); KPC *Kras*^G12D/fl^, n = 7, (n = 6 for BrdU)). Scale bar 200 µm. d) Quantification of BrdU and p53 positive cells, αSMA and picrosirius red positive area from PDAC of KPC *Kras*^G12D/+^ and KPC *Kras*^G12D/fl^ mice aged to clinical endpoint. (KPC *Kras*^G12D/+^, n = 7, (n = 6 for BrdU); KPC *Kras*^G12D/fl^, n = 7 (n = 6 for BrdU)). Data are mean ± s.e.m, *P* = 0.1970 (BrdU), \**P* = 0.0265 (p53), *P* = 0.8048 (αSMA), *P* = 0.2593 (picrosirius red), one-way Mann–Whitney U test.

**Supplementary Figure 4:**
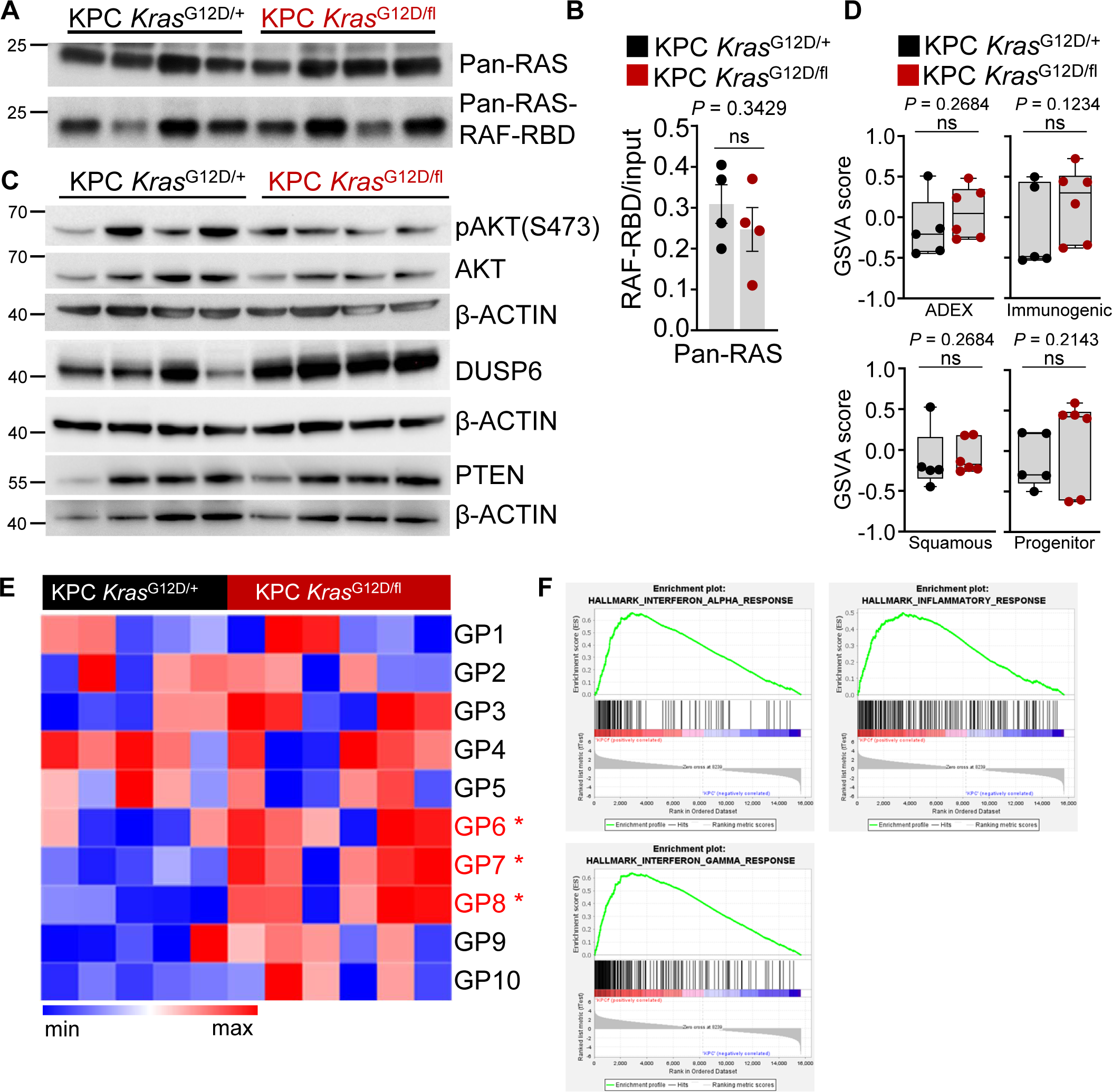
Loss of wild-type KRAS does not alter PI3K-AKT signalling. a) RAF-RBD agarose affinity purification assay of KPC *Kras*^G12D/+^ and KPC *Kras*^G12D/fl^ from PDAC tissue of four biological independent samples per condition. Pulldown of RAS-GTP with RAF-RBD agarose beads. Precipitates were immunoblotted using a pan-RAS antibody and input pan-RAS served as loading control. b) Bar graphs represents RAS-GTP activation levels quantified and normalized to pan-RAS loading control (from S4a)). N = 4 biological replicates per group, each lane represents snap frozen PDAC tissue lysate generated from individual mice from genotype indicated. Data represent mean ± s.e.m., *P* = 0.3429, one-way Mann–Whitney U test. c) Immunoblotting of pAKT (Ser473), AKT, DUSP6 and PTEN of KPC *Kras*^G12D/+^ and KPC *Kras*^G12D/fl^ of PDAC tissue lysates generated from mice at clinical endpoint. ß-ACTIN was used as loading control. Each lane represents an individual mouse of the indicated genotype. d) Boxplots of transcriptional pancreatic subtypes after Bailey classification of KPC *Kras*^G12D/+^ and KPC *Kras*^G12D/fl^ mice aged to clinical endpoint. Boxes depict interquartile range, central line indicates median and whiskers indicate minimum/maximum values (KPC *Kras*^G12D/+^,n= 5; KPC *Kras*^G12D/fl^,n = 6). e) Heatmap of gene programmes (GP) of Bailey classification of KPC *Kras*^G12D/+^ and KPC *Kras*^G12D/fl^ (KPC *Kras*^G12D/+^, n = 5; KPC *Kras*^G12D/fl^, n = 6). Significantly enriched GPs are highlighted in red. \**P* = 0.0411 (GP6, GP7), \**P* = 0.026 (GP8), one-way Mann–Whitney U test. f) Gene sets enriched in KPC *Kras*^G12D/fl^ tumours relative to KPC *Kras*^G12D/+^ tumours. Y-axis is enrichment score (ES) and x-axis are gene represented in gene sets.

**Supplementary Figure 5:**
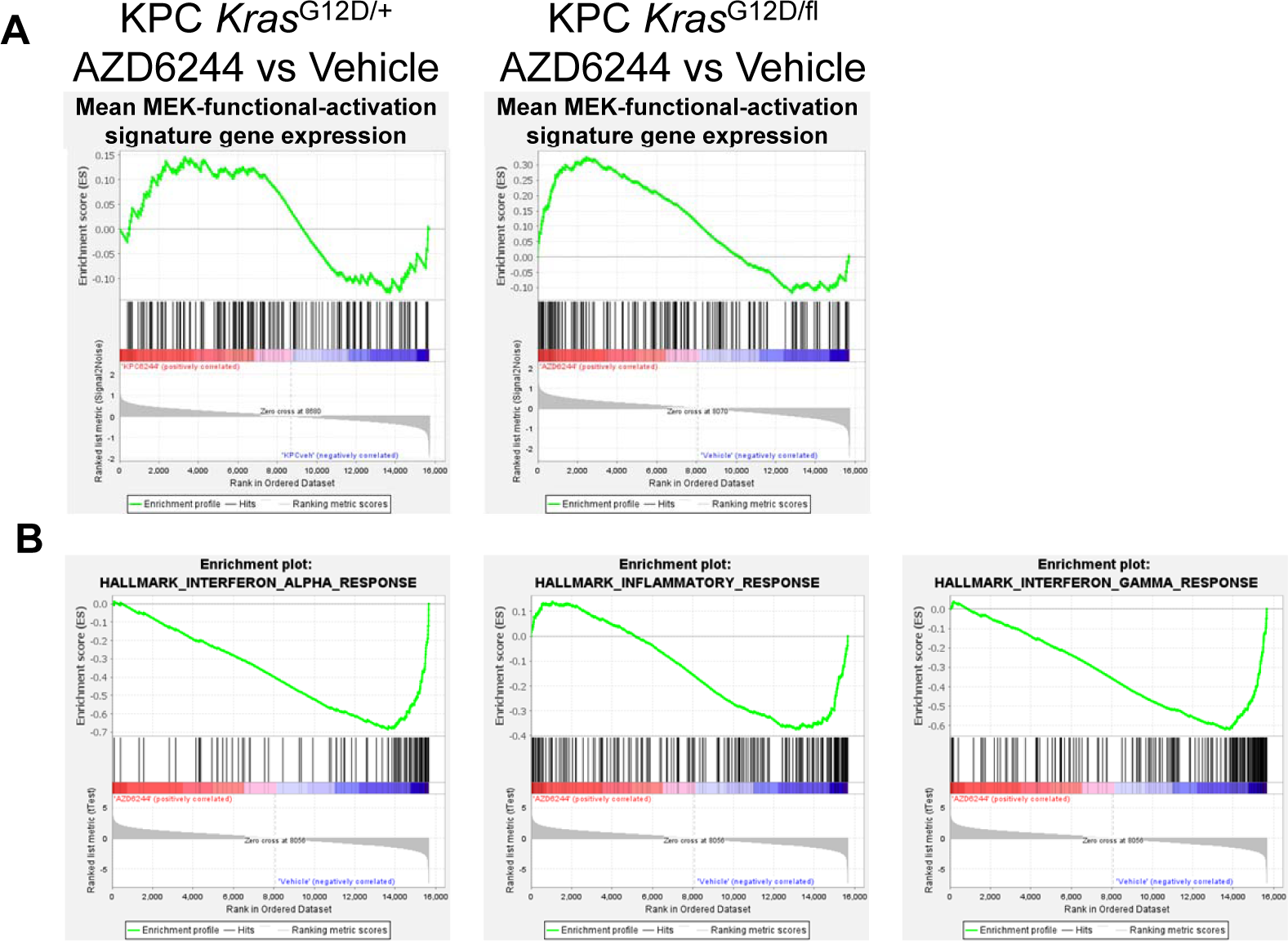
MEK1/2 inhibition in established tumour burden reverses enrichment of immune response related gene programmes in KPC *Kras*^G12D/fl^. a) Top: Gene expression signature for MEK-functional activation in KPC *Kras*^G12D/+^ (top) and *Kras*^G12D/fl^ (bottom) tumours treated with either vehicle or AZD6244. Y-axis is enrichment score (ES) and x-axis are gene represented in gene sets. b) Hallmark gene sets for immune regulatory pathways enriched in KPC *Kras*^G12D/fl^ tumours treated with AZD6244 relative to vehicle. Y-axis is enrichment score (ES) and x-axis are gene represented in gene sets.

**Supplementary Figure 6:**
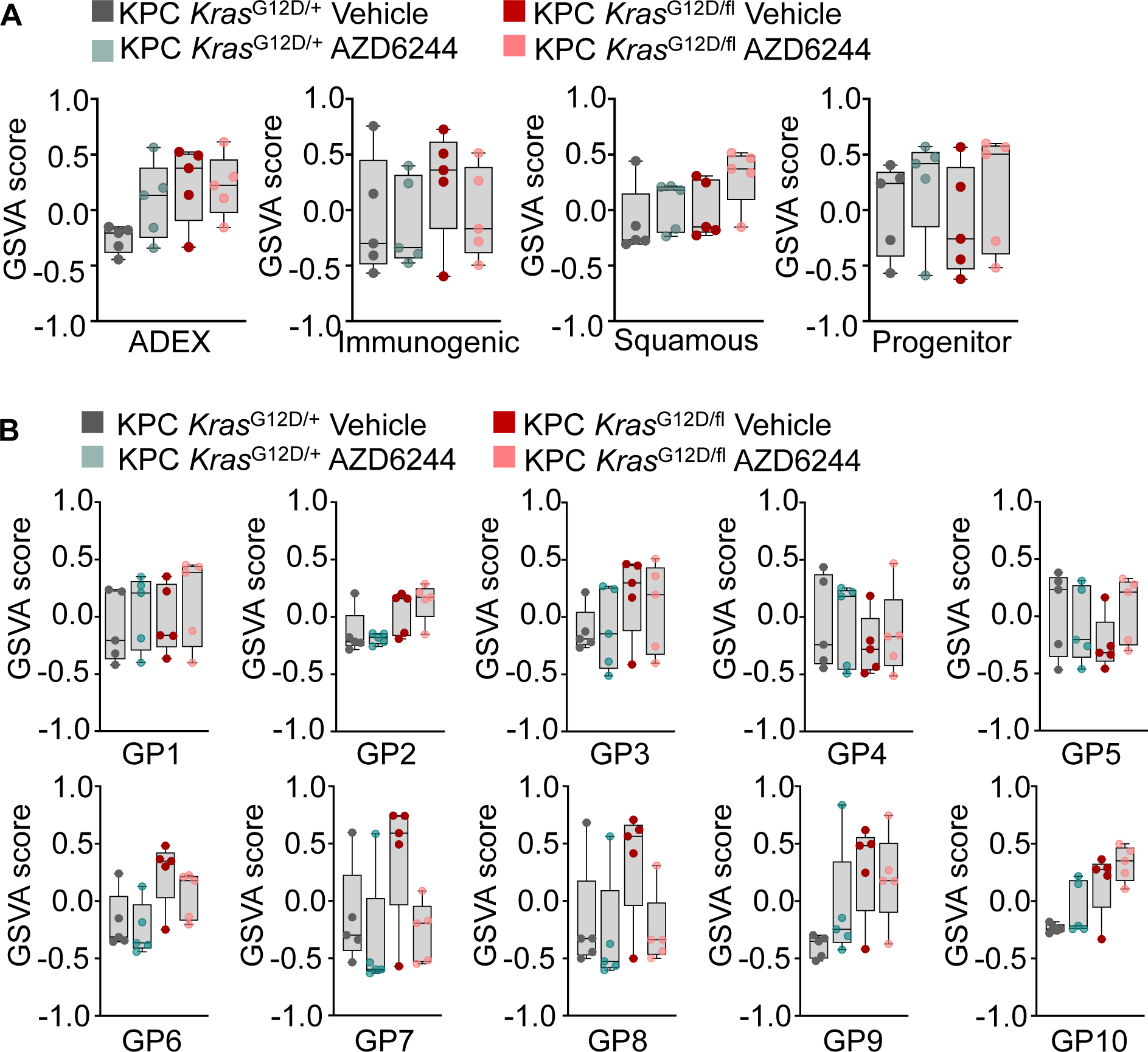
Loss of wild-type KRAS sensitizes KPC *Kras*^G12D/fl^ tumours to MEK1/2 inhibition. a) Boxplots of transcriptional pancreatic cancer subtypes after the Bailey classification for KPC *Kras*^G12D/+^ and KPC *Kras*^G12D/fl^ treated as indicated with either vehicle or AZD6244 from palpable tumour burden to clinical endpoint. Boxes depict interquartile range, central line indicates median and whiskers indicate minimum/maximum values (KPC *Kras*^G12D/+^ vehicle, n= 5; KPC *Kras*^G12D/+^ AZD6244, n = 5; KPC *Kras*^G12D/fl^ vehicle, n= 5; KPC *Kras*^G12D/fl^ AZD6244, n = 5). b) Boxplots of gene programmes (GP) of Bailey classification of KPC *Kras*^G12D/+^ and KPC *Kras*^G12D/fl^ treated as indicated with either vehicle or AZD6244 from palpable tumour burden to clinical endpoint. Boxes depict interquartile range, central line indicates median and whiskers indicate minimum/maximum values (KPC *Kras*^G12D/+^ vehicle, n= 5; KPC *Kras*^G12D/+^ AZD6244, n = 5; KPC *Kras*^G12D/fl^ vehicle, n= 5; KPC *Kras*^G12D/fl^ AZD6244, n = 5).

